# Emergent Collective Behavior Evolves More Rapidly Than Individual Behavior Among Ant Species

**DOI:** 10.1101/2024.03.26.586722

**Authors:** Grant Navid Doering, Matthew M. Prebus, Sachin Suresh, Jordan N. Greer, Reilly Bowden, Timothy A. Linksvayer

## Abstract

Emergence is a fundamental concept in biology and other disciplines, but whether emergent phenotypes evolve similarly to non-emergent phenotypes is unclear. The hypothesized process of *emergent evolution* posits that evolutionary change in collective behavior is irreducible to evolutionary change in the intrinsic behaviors of isolated individuals. As a result, collective behavior might evolve more rapidly and diversify more between populations compared to individual behavior. To test if collective behavior evolves emergently, we conducted a large comparative study using 22 ant species and gathered over 1,500 behavioral rhythm time series from hundreds of colonies and isolated individuals, totaling over 1.5 years of behavioral data. We show that analogous traits measured at individual and collective levels exhibit distinct evolutionary patterns. The estimated rates of phenotypic evolution for the rhythmicity of activity in ant colonies were faster than the evolutionary rates of the same behavior measured in isolated individual ants, and total variation across species in collective behavior was higher than variation in individual behavior. We hypothesize that more rapid evolution and higher variation is a general feature of emergent phenotypes relative to lower-level phenotypes across complex biological systems.

## Introduction

Collective behaviors such as the synchronization of firefly flashes (*1*), the collective movement of schools of fish (*2*) or flocks of birds (*3*), and the collective cognition of house-hunting ants (*4*) are conspicuous and widespread (*5*, *6*). Research on the mechanisms generating collective behavior emphasizes that simple rules governing the behavior of interacting individuals can lead to emergent features of a collective behavior (*7*). However, whether the emergence underlying trait expression has consequences for trait evolution remains unknown.

Because collective behaviors depend on the signaling and behavioral responses of interacting group members as well as the physiological and neural state of each individual group member (*6*, *8*), collective behavior might be more mechanistically complex and evolutionarily labile than the behavior of isolated individuals. Indeed, in theory, whenever trait expression depends on social interactions, the dynamics of trait evolution are expected to change, often including more rapid evolution and more phenotypic variation within and between populations (*9–11*). Nearly 100 years ago, WM Wheeler hypothesized that collective behavior may experience “emergent evolution”, whereby the evolution of a group-level trait may be irreducible to evolutionary changes in traits measured at a lower level (*12*, *13*). One consequence of such emergent evolution for collective behavior is that collective behavior might evolve faster and be more variable between species compared to the behavior of isolated individuals.

Empirical investigations into the evolution of collective behavior have been hindered by the difficulty of collecting a large sample of quantitative data on collective behavior from many species (*14*). Acorn ants (genera *Temnothorax* and *Leptothorax*) have strong potential as a model clade that can be leveraged to overcome this challenge. These ants are diverse, widespread, relatively easily collected, and well studied (*15*). Acorn ant colonies are small, often containing fewer than 200 workers, and colonies are readily cultured in artificial nests in the lab (*15*, *16*), where the behavior of whole colonies and isolated individuals can be quantified under controlled conditions (*4*, *17*).

Many traits, such as the cohesiveness of collective motion in bird flocks or the architecture of social insect nests are truly emergent in the sense that they exist and can be quantified at one level but not at a lower level (*18*). Other traits can be clearly defined and quantified at both the individual and collective level. These traits are also emergent as long as quantitative features at the collective level are not determined by quantitative features at the individual level (*19*). As a first test of emergent evolution across 22 ant species, we chose an exemplar case of the latter type of trait – biological rhythms of rest/activity – so that we could directly estimate and compare rates of phenotypic evolution at collective and individual levels. Rest/activity rhythms are ubiquitous across organisms, often impact fitness (*20*, *21*), and can be quantified using automated tracking methods (*22*, *23*). In this case, the null hypothesis is that evolutionary changes in features of activity rhythms at the group level are simply caused by evolutionary changes in the intrinsic activity rhythms of group members, so that interspecific variation for group-level activity rhythms should be strongly correlated with interspecific variation for individual-level activity rhythms. Alternatively, if ant activity rhythms exhibit an unambiguous case of emergent evolution, then 1) interspecific variation for colony-level activity rhythms should significantly exceed the interspecific variation of activity rhythms from isolated individuals and 2) the evolution of colony-level rhythm traits should be uncorrelated with the evolution of individual-level rhythm traits.

## Results

We filmed whole colonies and isolated individuals from 22 ant species, including 20 species of acorn ants (11 species of *Temnothorax* and 9 species of *Leptothorax*) and two outgroup species that also have small colony sizes and frequently inhabit acorns (*Tapinoma sessile* and *Myrmica punctiventris*). The rest/activity patterns of ant activity conform to a prominent ultradian rhythm (i.e., a biological rhythm with a periodicity shorter than 24hrs); workers inside of nests activate in synchronous bursts of locomotion (*17*, *22–24*) (Video S1), which has been shown to alleviate traffic congestion inside the nest cavity (*23*). We thus quantified both the rhythmicity (i.e., the regularity) and dominant period of oscillation for isolated individuals and entire colonies using wavelet analysis (*25*) as well as the average walking speed of lone workers in an open area (see supplementary methods section for details, Tables S1-3). To first test whether species differed for the measured individual and collective behaviors after statistically accounting for intraspecific variation, we used linear mixed models that included colony as a random factor. All of the measured behavioral traits except for individual rhythmicity differed between species (Figure 1a-c; Table S4). Our findings are consistent with the estimates of colony-level period in the handful of *Temnothorax* and *Leptothorax* species that have been previously studied (*17*, *22–24*, *26*).

**Fig. 1.**
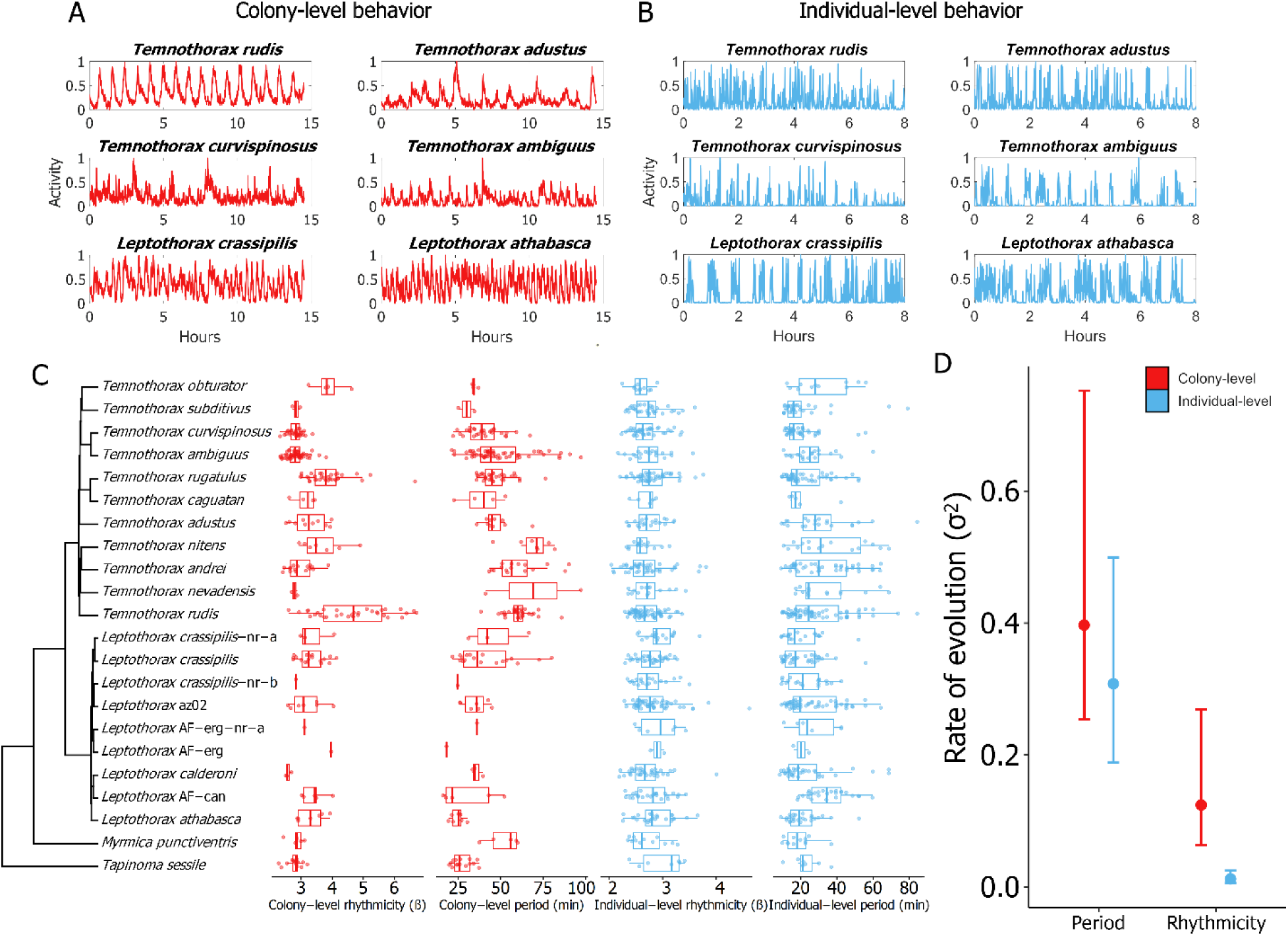
(a) Representative time series of collective activity from colonies of six different ant species. (b) Representative activity time series from isolated individuals of six different ant species. All of the visualized time series depict the raw data after being rescaled to fall between 0 and 1. (c) The phylogeny of species used in this study, with boxplots showing the data for each of our behavioral traits next to the corresponding species. Colony-level data is potted in red, and data from isolated individuals is plotted in blue. Each colony-level (red) data point represents the mean trait value for a unique colony. Each individual-level (blue) data point represents the data from a unique individual. (d) Estimates for the rates of evolution (σ^2^) for the matching collective and individual ultradian traits. The set of rate estimates for each trait was obtained using a bootstrapping approach (see supplementary methods). Dots represent rate medians and error bars depict the 95% confidence intervals.

To test whether the activity time series traits at the collective level exhibited distinct evolutionary patterns from the same traits measured at the individual level, we used phylogenetic comparative methods. First, we estimated the phylogenetic relationships among our study species based on a combination of existing DNA sequence data and new sequencing for this study (see supplementary methods). We then estimated rates of phenotypic evolution for each activity trait across the phylogeny. The estimated rates of evolution for collective rhythmicity were significantly higher than the rates for individual rhythmicity (Figure 1d; 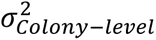 vs.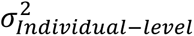 bootstrap p-value = 0.0001). In contrast, the rates of evolution for collective period and individual period were not significantly different from each other (Figure 1d; 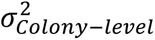 vs. 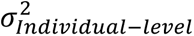 bootstrap p-value = 0.23). We tested whether the collective behaviors and individual behaviors were evolutionarily correlated using phylogenetically informed generalized least squares regression (PGLS) and we visualized the phylomorphospaces of each trait at the collective and individual level (Figure 2c, d). Analogous collective/individual trait pairs were not evolutionarily correlated (PGLS: rhythmicity – t-statistic = -0.01, p-value = 0.99; period – t-statistic = 0.29, p-value = 0.77). Performing a sensitivity analysis to account for the inherent intraspecific variation in our rhythm traits likewise supported this same conclusion (sensitivity PGLS: rhythmicity – p-value = 0.50; period – p-value = 0.39). We also tested whether the total multivariate phenotypic space (i.e. phenotypic disparity) (*27*) across our study species was different for the collective traits compared to individual traits. The estimated total phenotypic diversity of colony-level behaviors was significantly greater than the phenotypic diversity of individual-level behaviors (Figure 2a, b; collective-level disparity vs. individual-level disparity bootstrap p-value = 0.0001). Lastly, we tested whether species’ colony-level ultradian rhythm traits might be determined by an influential minority of individuals within colonies (*28*) by performing a “disassembly experiment” where we extracted all individuals from three arbitrarily selected colonies (one colony of *Temnothorax obturator*, *Temnothorax rudis*, and *Leptothorax crassipilis*) and measured every ant in isolation concurrently (see supplementary results).

**Fig. 2.**
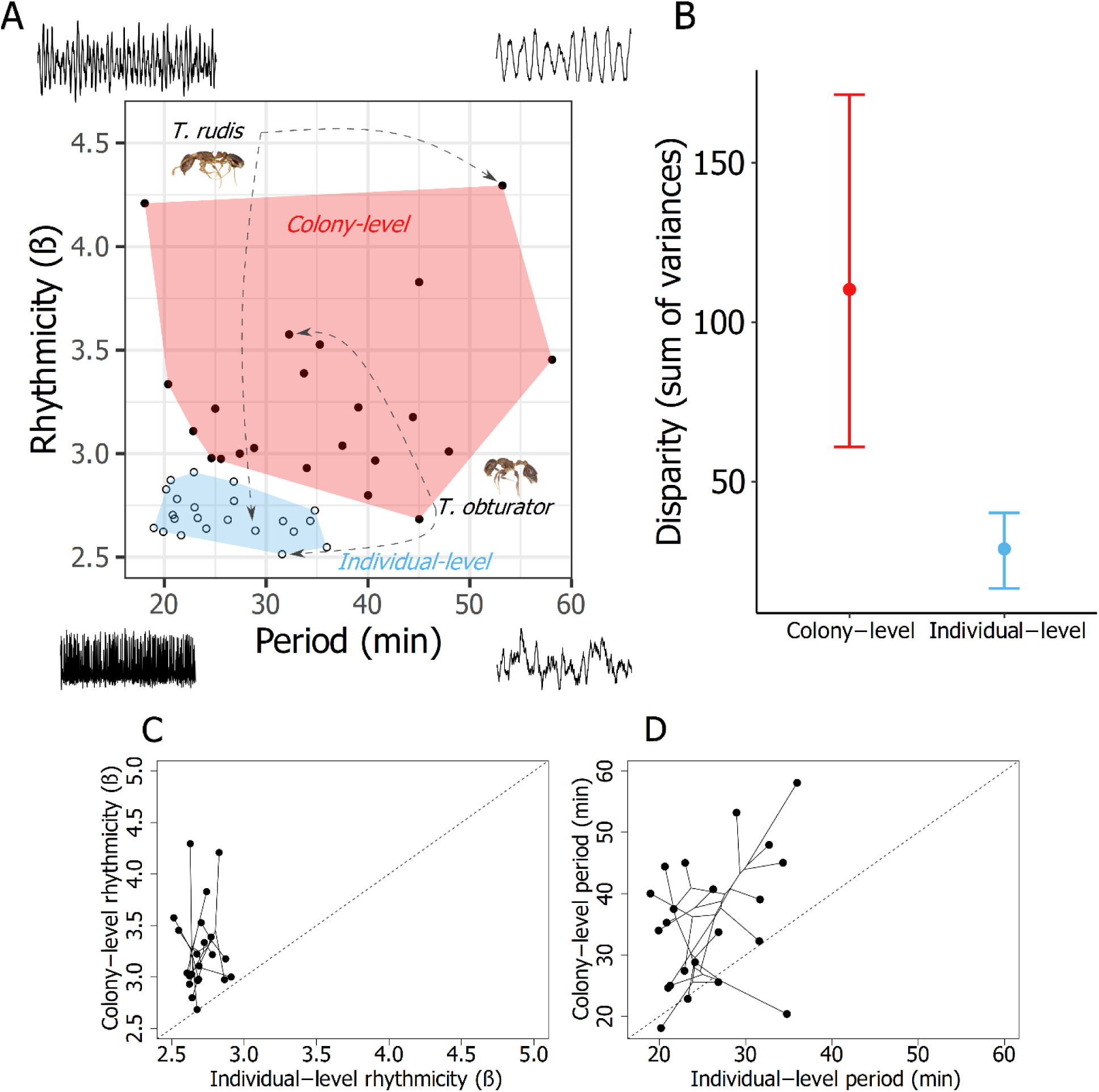
(a) Behavioral phenospace of ant ultradian rhythms across 22 different species. Each datapoint represents a different species. Solid points falling within the pink polygon correspond to species’ colony-level rhythms, and open circles within the blue polygon correspond to species’ individual-level rhythms. The individual-level and colony-level datapoints associated with *Temnothorax rudis* and *Temnothorax obturator* are provided as examples. Specimen images are to scale and were extracted from images from www.antweb.org (*T. rudis*: casent0005689, *T. obturator*: casent0104756) Each corner of the phenospace is ornamented by a synthetic time series generated by a noise-driven FitzHugh-Nagumo model (see supplementary methods) that qualitatively illustrates the features of time series from their respective region of the phenospace. (b) comparison of the phenotypic disparity (measured as the sum of variance) between species’ colony-level and individual-level behavior spaces. The sum of variances were calculated by bootstrapping the phenospace data. Dots represent medians and error bars depict the 95% confidence intervals. (c) Behavioral phylomorphospaces for individual-level vs. colony-level period and (d) individual-level vs. colony-level rhythmicity. The dotted lines show what a 1:1 relationship between the variables would be.

When considering both rhythmicity and period together with our multivariate phenospace analysis, ant ultradian rhythms show evidence of emergent evolution because the total phenotypic space of colony-level ultradian behavior is larger and occupies a different region of phenospace than individual-level behavior (Figure 2a, b). When considering each trait separately, the rhythmicity of ant activity meets our criteria for unambiguous emergent evolution: 1) colony-level rhythmicity evolved more rapidly than individual-level rhythmicity and 2) colony-level rhythmicity and individual-level rhythmicity were not evolutionarily correlated. In contrast, the period of ant activity rhythms does not meet both of our criteria because the rate of evolution for colony-level period was not significantly higher than the rate for individual-level period.

## Discussion

Nearly a century after the concept was originally proposed (*29*), we have uncovered empirical evidence for the emergent evolution of collective behavior. Using automated behavioral tracking to characterize the behavior of both isolated individuals and entire colonies across 22 species, we found that colony-level ultradian rhythmicity evolved more rapidly than the same trait expressed by isolated individuals. Consequently, the total phenotypic divergence of ultradian rhythm behavior across our study species was much larger at the colony level than at the individual level.

The collective behavior of groups fundamentally depends on social interactions among group members (*6*), but it is unknown whether the evolution of collective behavior is primarily achieved by modifying social interaction rules or by modifying the intrinsic behavior of individuals. Social factors are known to strongly affect behavioral rhythms in ants. For example, the presence of brood items can drastically alter the expression of both circadian and ultradian rhythms in ants (*30*, *31*), removing a colony’s queen dampens colony synchrony (*24*), inactive ants inside the nest can be roused by physical contact with moving ants (*17*, *32*), and increasing the number of ants in a small group (i.e., 1, 3, 7, or 15 ants) causes more rhythmic collective activity rhythms (*33*). Our discovery of faster evolution of a collective trait compared to the *same* trait measured in individuals suggests that interspecific differences in response to these social factors – i.e. interspecific differences in social interaction rules -- exert substantial influence on the evolution of emergent collective behaviors. The interspecific differences in emergent rhythmicity expressed by groups of individuals in a colony context therefore appears irreducible to interspecific variation in an intrinsic rhythmicity that is independent of social context.

Our findings likely have broad relevance to the phenotypic evolution of collective behavior in general as well as the specific case of the evolution of behavioral rhythm synchronization. Various forms of synchronized behavior have repeatedly and independently evolved across organisms (*1*, *5*, *34*, *35*). Both within and across lineages that exhibit synchronized behaviors, there is considerable variation in the properties of synchronized behavior. For instance, different species of fireflies vary in the consistency of their synchronized flashing, with many species lacking synchronized displays altogether (*36*, *37*). Similarly, synchronized waving displays in fiddler crabs also vary widely between species for degree of synchrony (*38*, *39*). If our finding that colony-level rhythmicity evolves faster than individual-level rhythmicity in ants is generalizable, then social interaction/coupling rules might be more important than individual rhythm characteristics in causing the evolution of species-level differences in biological synchronization phenomena. We therefore predict that species should tend to have stronger interspecific variation in the properties of their synchronized behavioral rhythms compared to the properties of their behavioral rhythms when expressed in social isolation. A recent study on the biparental synchronization of egg incubation rhythms in shorebirds is consistent with this prediction; the incubation rhythms of synchronized shorebird parents observed in the field are highly variable across species, but this interspecific rhythm variation appears to be absent when captive birds are tested in social isolation (*40*). The reported discrepancy between the rhythmic behavior of synchronized wild birds compared to captive, isolated birds is expected in light of our findings and the predictions of emergent evolution.

We predict that emergent evolution should also occur much more generally across all emergent traits. This includes most collective behaviors as well as situations where a collective trait has no directly analogous individual trait associated with it, such as the cohesiveness of collective motion in bird flocks or nest structures built by the collective action of groups of social insects. Interspecific differences in nest architecture across multiple social lineages are well documented (*41*, *42*), and the production of nest structures can be achieved by interacting individuals following simple behavioral rules that depend on social context (*41*, *43*, *44*). If emergent evolution is widespread, then rapid evolution of large interspecific differences in emergent traits like nest structure may be achieved through the evolution of small changes in the social interaction rules followed by individuals.

Early theorists of emergent evolution also emphasized its potential for generating diversity in novel emergent traits and for underlying major evolutionary transitions (*29*), such as the evolution of eusociality (*12*). If true, we would expect that relatively small phenotypic changes in the traits of lower-level units (e.g., isolated cells or individuals), together with small changes in traits regulating social interactions, could readily generate relatively large phenotypic changes in the higher-level emergent phenotype, potentially underlying major transitions in biological organization. Available comparative genomic studies seeking to identify genetic changes underlying the evolution of multicellularity (*45–47*) and eusociality (*48–50*) emphasize such relatively small genetic changes, consistent with emergent evolution. Further study across a wide range of emergent traits and systems will be necessary to elucidate to what degree emergent evolution has contributed to generating the phenotypic diversity of life.

## Methods

### Colony information

The colonies used for this study were collected from multiple locations, all within the United States (Table S1). Colonies were maintained in plastic nestboxes (11×11×3 cm), and each colony was made to inhabit an artificial nest with a standardized design. These nests consisted of a 1.5mm thick chipboard slat (50×75 mm) with a rectangular hole (29×44 mm) that acted as a cavity the ants could inhabit. A 1.5mm wide and 4mm long slit was cut into each nest to serve as an entrance to the nesting cavity. Two glass microscope slides were used for each nest’s floor and roof, respectively. The larger body size of *M. punctiventris* necessitated a larger variant of this design; this variant consisted of a 3D printed nest (75×50×4 mm) with a rectangular nesting cavity (28×60 mm), a 9 mm wide entrance, and a microscope slide nest roof. Colonies were fed weekly with frozen *Drosophila melanogaster* and had *ad libitum* access to water as well as *ad libitum* access to a sucrose solution (Sunburst Ant Nectar - byFormica Ant Products). Prior to conducting the disassembly experiment, colonies were fed with *D. melanogaster* and a protein-free variant of the Dussutour-Simpson synthetic diet (*51*).

### Phylogenetic tree construction

We sampled 25 taxa for our phylogenetic tree, including all 22 species used in this study, with *Eciton burchelli* used as an outgroup. Samples of *Leptothorax* AF-can, *Leptothorax* AF-erg, *Leptothorax crassipilis* nr-a were all taken from the colonies used in this study. The remaining samples are conspecific with the study samples but were not taken directly from the colonies. Additionally, we incorporated nine samples from previously published datasets (*15*, *52*, *53*). In total we prepared 16 new samples for this project (see Table S5).

We extracted DNA, prepared genomic libraries, and performed targeted enrichment of ultraconserved elements (UCEs). For most of our samples, we extracted DNA nondestructively from adult worker ants by using a flame-sterilized size 2 stainless steel insect pin to pierce the cuticle of the head, mesosoma, and gaster on the right side of the specimens. For one pupal worker sample, we ground the specimen with a pestle in a 1.5 mL tube prior to extraction. In both cases, we then used a DNeasy Blood and Tissue Kit (Qiagen, Inc., Hilden, Germany) to extract DNA, following the manufacturer’s protocols. We verified DNA extract concentration using a Qubit 3.0 Fluorometer (Invitrogen, Waltham, MA, U.S.A.). We input up to 50 ng of DNA, sheared to a target fragment size of 400–600 bp with a QSonica Q800R sonicator (Qsonica, Newtown, CT, U.S.A.), into a genomic DNA library preparation protocol (KAPA HyperPrep Kit, KAPA Biosystems, Wilmington, MA, U.S.A.). For targeted enrichment of UCEs, we followed the protocol of Faircloth et al. (*54*) as modified by Branstetter et al. (*52*) using a unique combination of iTru barcoding adapters ((*55*); BadDNA, Athens, GA, U.S.A.) for each sample. We performed enrichments on pooled, barcoded libraries using the catalog version of the Hym 2.5Kv2A ant-specific RNA probes ((*52*); Arbor Biosciences, Ann Arbor, MI, U.S.A.), which targets 2,524 UCE loci in the Formicidae. We followed the library enrichment procedures for the probe kit, using custom adapter blockers instead of the standard blockers ((*55*); BadDNA, Athens, GA, U.S.A.), and left enriched DNA bound to the streptavidin beads during PCR, as described in Faircloth et al. (*54*). Following post-enrichment PCR, we purified the resulting pools using SpeedBead magnetic carboxylate beads ((*56*); Sigma-Aldrich, St. Louis, MO, U.S.A.) and adjusted their volume to 22 μL. We verified enrichment success and measured size-adjusted DNA concentrations of each pool with qPCR using a SYBR-FASTqPCR kit (Kapa Biosystems, Wilmington, MA, U.S.A.) and a Bio-Rad CFX96 RT-PCR thermal cycler (Bio-Rad Laboratories, Hercules, CA, U.S.A.), subsequently combining all pools into an equimolar final pool. We sequenced the final pool in one lane at Novogene (Sacramento, CA, U.S.A.) on Illumina HiSeq 150 cycle Paired-End Sequencing v4 runs (Illumina, San Diego, CA, U.S.A.), along with other enriched libraries for unrelated projects. Raw sequence reads can be found on the NCBI sequence read archive (BioProject PRJNA1060740).

We followed the standard PHYLUCE protocol for processing UCEs in preparation for phylogenomic analysis, aligning the monolithic unaligned FASTA file with the phyluce_align_seqcap_align command, using MAFFT (*57*) as the aligner (--aligner mafft) and opting not to edge-trim the alignment (–no-trim). We trimmed the resulting alignments with the phyluce_align_get_gblocks_trimmed_alignments_from_untrimmed command in PHYLUCE, which uses GBlocks ver. 0.91b (*58*) using the following settings: b1 0.5, b2 0.5, b3 12, b4 7. After removing UCE locus information from taxon labels using the command phyluce_align_remove_locus_name_from_nexus_lines, we examined the alignment statistics using the command phyluce_align_get_align_summary_data and generated a dataset in which each locus contains a minimum of 85% of all taxa using the command phyluce_align_get_only_loci_with_min_taxa. Following sequencing, assembly, and *in silico* extraction of UCE loci, we recovered an average coverage depth of 42.6x (range: 9.9-77.7x) and a mean contig length of 1226 bp (range: 59-8244 bp). After alignment, trimming, and filtering of UCE loci, the resulting matrix was 2.5 Mbp long and contained 2249 UCE loci, which had a mean length of 1114 bp (range: 320-2087 bp). The final matrix contained 15.6% gaps or missing data. For additional assembly statistics, see Table S6.

Because the assumption that the evolutionary rates of sequence data are homogenous is often violated in empirical data (*59*), we partitioned our UCE loci into sets of similarly evolving sites. To achieve this, we used the command phyluce_align_format_nexus_files_for_raxml which concatenates loci into a single alignment and generates a partition file for input into the SWSC-EN method (*60*). We used the resulting datablocks as input for partitioning in IQTREE v2.1.2 (*61*), using the command -m TESTNEWMERGEONLY. We set the substitution model to ‘general time reversible’ (-mset GTR). Because the combination of gamma and proportion of invariable sites (+I+G) has been demonstrated to result in anomalies in likelihood estimation (*62*, *63*), we set the rate heterogeneity models to a subset that includes everything except the combination of gamma and proportion of invariable sites (-mrate E, I, G). We set the search algorithm to -rclusterf 10. We used the resulting partitioned dataset as input for maximum likelihood tree inference in IQTREE using 1000 ultrafast bootstrap replicates (-bb 1000).

We used MCMCTREE in PAML v4.9e (*64*, *65*) with the approximate likelihood calculation to decrease the overall analysis time (*66*), to estimate a chronogram. We used one fossil calibration point and four secondary calibrations from the literature on our tree (see Table S7). We set the clock to the independent rates model (clock = 2) and used the HKY85 substitution model (model = 4). We set the gamma prior for overall substitution rate for genes (rgene gamma) to 1 5 using the results from an initial clock analysis in baseml. Finally, we set the number of samples to 10,000 with a sample frequency of 5,000 and a burnin of 5,000,000. We ran two analyses in parallel to confirm that results were stable and checked for convergence using the program Tracer (*67*).

### Filming and automated tracking of colony & individual activity

Each filming session for colonies lasted for 14.5 hours. We recorded colonies during these 14.5-hour sessions using DSLR cameras (Panasonic LUMIX) that were programmed to capture a still image of the colony’s nest every 30 seconds. The cameras were also programmed to begin shooting at 9:10 each morning. We aimed to record each colony on at least two separate occasions within 5 months of field collection to minimize any impacts of captivity on their behavior (*68*). To record the activity of isolated individuals, we removed workers that were walking around in colony nestboxes and placed them in 1.5ml microcentrifuge tubes. A small piece of molding clay (Gorilla Glue Company, Cincinnati, Ohio, USA) was packed into the cap of each tube to prevent ants from entering the top region of the tube, which would have obscured them from the camera’s view. The bottom of each tube was also filled with a piece of moist cotton, as we were initially concerned about the potential for desiccation. These tubes were then arranged in a grid underneath our DSLR cameras and filmed for 8.5 hours. Just as in our recordings of colonies, images of isolated individuals were obtained at 30 second intervals. All recorded individuals were eventually returned to their source colonies, but they were kept in their respective microcentrifuge tubes until we had finished recording individuals from their original source colonies. This ensured that no single individual ant would be recorded more than once, and we were therefore able to avoid any potential pseudoreplication. As is the case for at least one other ant genus (*69*), acorn ant workers inside the nest do not appear to possess endogenous circadian rhythms (*17*, *22*, *24*, *70*); their ultradian rhythms are stable under constant ambient conditions. We therefore conducted all of our recordings under constant temperature and light conditions.

The time series of colonies’ collective activity were automatically obtained using an image segmentation approach that is described in detail in previous work (*24*, *33*). Briefly, the nest area of a focal colony was first selected as a region-of-interest, and then adaptive image thresholding was applied to each frame of the focal colony’s 14.5-hour long image sequence. The proportion of pixels that change between frames is used to estimate the proportion of individuals that have moved within the nest between successive frames of the image sequence. This technique provides an accurate estimate of the approximate proportion of individuals in a colony that have moved during a given interval of time (every 30 seconds in our case); it yields results that are comparable to those provided by using automated tracking on colonies that are fully marked with computer readable tags. Using a marker-less, threshold-based approach allowed us to collect data from many more colonies than would be feasible with the more detailed tag-based methods, and also allowed us to avoid the potential negative effects of tagging on the ants’ other behaviors (*71*). When we recorded colonies prior to our disassembly experiment, our DSLR cameras were unavailable, so we filmed colonies using continuous video at 720p using camcorders (Canon VIXIA). Frames were extracted from these videos at 30sec intervals so that the sampling interval of their calculated activity time series would match the rest of the time series in the study. Our filming setup combined with the lower resolution of these video frames compared to our DLSR images meant that we needed to use an alternative approach to measuring the collective activity in these recordings; a much higher level of image noise made the image thresholding method unsuitable. To overcome this issue, we used optical flow, which prior work has successfully used to track the motion of *Leptothorax* (*23*). This method identifies changes in pixel brightness values between frames of a video or image sequence to estimate the speed of moving objects (see miscellaneous methodological details section below for additional information).

All isolated worker ants were filmed with a camcorder (Canon VIXIA) on an 8.5×11in sheet of pink paper to measure their walking speed in an open field assay. These open field assays lasted until either the ant walked off the paper or approximately 1min had elapsed since the beginning of the test. Sheets were replaced for each ant. Frames were extracted from these videos of these open field assays at a rate of 5 fps. The position of the ant in each frame was automatically detected using image thresholding, and we calculated the Euclidian displacement of the ants’ centroid between successive frames. To account for the fact that larger ants will tend to naturally walk faster than smaller ants (*72*), we standardized our measurement of the instantaneous walking speed of each ant by dividing the ant’s speed (originally in pixels/second) by the average body size of the ant (measured in pixels), resulting in velocity units of body-lengths/sec. The mean moving speed for each ant’s assay was then determined by taking the average of the velocity time series when the ant was in motion, which was defined as whenever the ant’s instantaneous velocity exceeded 0.5 body-lengths/s. After experiencing her 1min walking assay, each ant was transferred to a 1.5ml microcentrifuge tube for the individual-level rhythm recordings. Just as was done for the open field assay, isolated individuals in microcentrifuge tubes were tracked by automatically identifying their coordinates in each frame of their respective recording and calculating their Euclidean displacement between successive frames (i.e., every 30 seconds).

Isolated individuals in our disassembly experiment were kept in circular plastic arenas (diameter: 8 mm) that were topped with a glass microscope slide. These arenas, being smaller than the microcentrifuge tubes used for the main experiment, allowed us to comfortably film all of the isolated individuals from each colony concurrently. Like the colony videos from the disassembly experiment, we tracked the ultradian rhythms of the isolated ants using optical flow due to similar issues stemming from image noise in the camcorders. In addition to the three species presented in our study (*T. obturator*, *T. rudis*, and *L. crassipilis*), we also disassembled and filmed the individuals from the colony of a fourth species (*T. ambiguus*), but due to an error setting up the camera, the resulting footage was too overexposed, and the *T. ambiguus* individuals (which naturally have a light integument) were neither visible nor trackable in the video.

### Quantifying tempo traits

We calculated two simple metrics (rhythmicity, and dominant period) to characterize essential features of each time series in our study. These two metrics were obtained using wavelet analysis, a technique that has seen successful application in several ant behavior studies (*17*, *23*, *24*, *73*). Both of the time series metrics were calculated by creating wavelet periodograms for each time series, which were computed by applying the MATLAB wavelet-transform function *cwt*. We summed all of the wavelet magnitudes associated with each frequency present in the wavelet transform’s output. After converting frequency to period, this gave us the total power associated with different oscillation periods in a given time series.

The dominant period of an ant/colony’s time series expresses the timescale of the underlying oscillatory behavior. It can be thought of as the typical interval between bursts of activity for a given time series. We determined the dominant oscillation period for a given time series by finding the highest peak in the wavelet periodogram and determining the oscillation period associated with this peak. The rhythmicity metric conveys how predictable and consistent the activity oscillations are. To quantify the rhythmicity of time series in our dataset, we calculated a coherence factor (*β*), a widely used approach which can be thought of as a signal-to-noise ratio (*74*, *75*). We defined this metric as the ratio of the height of the periodogram’s tallest peak (i.e., the dominant period) to the mean of the wavelet magnitude values in the periodogram. Larger values of *β* indicate that the activity oscillations are more predictable, prominent, and stable.

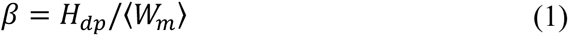

Prior to calculating these two time series metrics, we smoothed all of the time series with a Gaussian-weighted moving average with a 15-point window. This smoothing is essential as previous work has established that high frequency noise in the time series can cause the wavelet analysis to report inaccurate results unless this amount of smoothing is applied (*17*).

### Data analysis and model simulation

To test for the presence of interspecific differences in our five behavioral traits, we built linear mixed-effects models using the R package nlme. Species ID was included as a fixed effect, colony ID (or colony of origin in the case of individual ants) was included as a random effect, and the response variable was one of the five behavioral traits. For our phylogenetic signal, PGLS, and disparity analyses, we required a single taxa-specific value for each trait. To account for intraspecific variation and the fact that most colonies were filmed twice for the collective traits, we used the coefficient estimate associated with each species in the LME models to get a tip-value for each species. We performed our analyses of phenotypic disparity using the R package dispRity (*27*). Phylomorphospace plots were generated with the phytools package in R. Our PGLS analyses were performed in R with the packages phytools and sensiPhy. The sensitivity tests of PGLS and phylogenetic signal in the sensiPhy package all used 10,000 sampling iterations. We estimated the rates of evolution for each trait using a bootstrapping approach. First, we randomly sampled (with replacement) the datasets of both individual and collective activity to create 10,000 new datasets of the same size as the original data. For each resampled dataset, we then computed rate estimates for each trait by fitting a continuous Brownian motion model using the fitContinuous function in the R package geiger. We tested if the resulting rate estimates of the two collective traits were significantly different from their individual counterparts by calculating the difference between the rates (e.g., 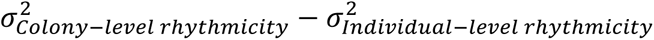) and observing what proportion of the rate differences exceeded the null expectation (i.e., a rate difference of zero). We standardized our behavioral metrics prior to estimating the evolutionary rates; for each trait, we divided every species’ tip-value by the mean of all tip-values for that trait. Because the individual-level time series were shorter than the colony-level time series, all our analyses that compared colony-level vs. individual-level traits were done using truncated colony-level time series so that we could compare time series of equal length. All statistical analyses were conducted using R version 4.0.3 (https://www.r-project.org).

## Data availability

All data (including all time series) and analysis code used for this study are available on GitHub (https://github.com/naviddio/Comparative_tempo) and will be archived on Zenodo upon acceptance. Sequence data are available on the NCBI sequence read archive (BioProject PRJNA1060740).

## Acknowledgements

We thank the Wisconsin Department of Natural Resources, the city government of Austin (TX), the city government of Bloomington (IN), the city government of Portland (OR), the UC Sagehen Creek Field Station (Reserve DOI: 10.21973/N33M26), the UC James San Jacinto Mountains Reserve (Reserve DOI: 10.21973/N3KQ0T), the Brackenridge Field Laboratory (Austin, TX) of UT-Austin, the Forest Preserves of Cook County (IL), the Ridges Sanctuary (Door County, WI), and the Pack Experimental Forest (Eatonville, WA) of the University of Washington for allowing us to collect ants on their properties. We also thank Christian Rabeling and members of the Social Insect Research Group at Arizona State University for feedback on this project.

## Author contributions

GND and TAL conceived of and designed the research; GND, SS, JNG, and RB performed the research; MMP conducted the genetic sequencing and built the phylogeny; GND and TAL analyzed the data; GND wrote the initial draft of the paper. GND, TAL, SS, and MMP contributed to the editing of the paper.

