## Supplementary material for "Emergent Collective Behavior Evolves More Rapidly Than Individual Behavior Among Ant Species"

### **Supplementary methods**

*Miscellaneous methodological details*

The optical flow method used for our disassembly experiment has been previously implemented for studying ant activity (*1*). Our use of optical flow relied on the Farneback technique. Greater details on this method can be found in other sources (see refs. (*1*, *2*)), but the basic principle behind this technique is that changes in the brightness values of pixels in successive frames of an image sequence are used to estimate the motion of objects in the frames. An optical flow vector is calculated for each pixel in a pair of frames, and a given optical flow vector conveys the approximate direction and velocity (pixels/frame) of the object for that pixel. Larger average optical flow magnitudes can thus be used as a proxy for the proportion of ants in the nest that are active in the case of colony recordings or individual walking speed in the case of isolated individuals.

Five of the colonies analyzed in this study were chimeric. Specifically, two out of the 29 *Temnothorax curvispinosus* recorded for our study contained the dulotic social parasite *Temnothorax americanus* and another two of the colonies contained the workerless inquiline *Temnothorax minutissimus*. These colonies were classified as *T. curvispinosus* in our analyses since these parasites were only a small minority of each nest's population. Additionally, one of the colonies classified as *Leptothorax athabasca* contained a minority of workers and larvae of *Leptothorax calderoni*. The two *Leptothorax* AF-erg ants used for the individual-level recordings were taken from a small colony that was collected in 2021. At the time of our study, this colony had declined and only consisted of these two ants. The large *L*. AF-erg colony that was used for our colony-level activity recordings did not have any foragers in their nestbox on the occasions of our individual-level recording sessions, so we resorted to using the two *L*. AF-erg individuals from this other colony fragment for our individual-level assays.

In our recordings of isolated individuals, ants would occasionally fail to be detected by our tracking code. We therefore only included individual-level time series in our analysis if the focal individual was detected in at least 70% of the frames from the recording. Because instances of failed detection generally occur when the ant is resting and occluded by part of the observation arena, we followed the protocol used in previous work and assigned a value of zero prior to the missing data-points in the activity time series that were used for the analyses prior to the application of the Gaussian smoothing (*3*).

We took a colony-level recording of a *Temnothorax emmae* nest, but workers of this colony appeared injured, and no workers were used in individual-level tests, so we did not include it in our study. A single worker of *Temnothorax wardi* was also recorded during our individual-level tests, but we did not have a colony to record (only three workers), so we could not include this species in our analyses. Also, tracking issues prevented the velocity data from one *Leptothorax athabasca* worker from being used in the open field velocity analysis, so there were 576-2=574 total workers whose velocity data was analyzed from the open field assay. There were tracking issues with one of the recordings of colony WRT2 (a *T. nitens* colony) and one of the recordings of colony BFLT2 (*T. obturator*), so these two colony-level recordings needed to be excluded. Two of the *Leptothorax Athabasca* individuals recorded for the open field assay were pseudogynes, which are unmated queens that behave like workers (*4*). Pseudogynes are not uncommon in colonies of *Leptothorax* (*5*).

In our disassembly experiment, a small proportion of ants from the *Temnothorax rudis* colony as well as the *Temnothorax obturator* colony escaped their arenas during filming and needed to be returned to their respective nestboxes. Five workers escaped (out of 70 ants total) in the *T. rudis* recordings, one worker escaped (out of 18 ants total) in the *T. obturator* recordings, and one worker escaped (out of 16 ants total) in the *L. crassipilis* recordings. These escaped workers were thus excluded from the analysis of the disassembly experiment.

The four illustrative time series that appeared in the corners of our behavioral phenospace (main text Figure 2A) were generated by numerically simulating a FitzHugh-Nagumo model. The FitzHugh-Nagumo model was originally designed to describe the behavior of neural oscillations but is often used across disciplines as a generic formulation of an excitable system (*6–8*). The model consists of two differential equations; our noise-excited variant of this model is the same as the form presented by Pikovsky and Kurths (*9*).

$$\epsilon \frac{dx}{dt} = x - \frac{1}{3}x^3 - y \quad (1)$$

$$\frac{dy}{dt} = x + a + D\xi(t) \quad (2)$$

In equations 2 and 3, the variables $x$ and $y$ represent the behavioral dynamics of the oscillator (membrane voltage and neuron recovery in the model's original context). The $\xi$ term represents the application of Gaussian noise, and $D$ is the amplitude of the added noise. The terms $a$ and $\epsilon$ represent adjustable parameters of the systems. The example time series used in Figure 2A from the main text plot the $y$ variable. We ran simulations of this model using the deSolve package in R.

The MATLAB function sort_nat was used to ensure that our image sequences were correctly sorted in chronological order for our image analyses. This function was downloaded from the MathWorks File Exchange: (https://www.mathworks.com/matlabcentral/fileexchange/10959-sort_nat-natural-order-sort).

### **Supplementary results**

*Colony size analysis*

We assessed the effect of colony size by looking at the four species with the greatest number of colonies in our dataset (*T. ambiguus*, *T. curvispinosus*, *T. rudis*, and *T. rugatulus*). For each species, we fit an LME model with rhythmicity or period as the response variable, colony size (i.e., approximate number of adult ants inside the nest) as the fixed effect, and colony ID as a random effect. Colony size was not correlated with either of our colony-level traits in any of the four species (Figure S2a, b; Table S8). A PGLS analysis likewise found no correlation for colony-level period vs. colony size (PGLS: t-statistic = 1.23, p-value = 0.23) nor colony-level rhythmicity vs. colony size (PGLS: t-statistic = 1.93, p-value = 0.0681).

*Disassembly experiment*

Because our tests of isolated individuals relied on only a subset of ants from a given colony, it is possible that colony-level ultradian behavioral activity rhythms are determined by the average behavior of a specific behavioral caste (e.g., nurse workers) or a small number of especially influential ants within the colony (e.g., keystone individuals) (*10*). Our disassembly experiment tested this by isolating every ant from a colony in separate arenas and measuring their activity rhythms. We compared the distribution of individual-level time series activity traits in a colony with the corresponding colony-level values, which we obtained using automated analysis of the recording of the full, intact colony taken two days prior to the disassembly experiment.

Our results support the hypothesis that group-level characteristics of ants' collective ultradian rhythms (specifically the colony-level rhythmicity trait) are strongly contingent on the social environment of individual ants, i.e. that colony-level rhythmicity is an emergent trait. The colony-level rhythmicity values for the *T. rudis* and *T. obturator* colonies fell well outside of the distributions of individual-level rhythmicity values of isolated workers from these two colonies (Figure S3). In *T. rudis*, 98% of the individual rhythmicity values were lower than the colony value, and 100% of the individual rhythmicity values were lower than the colony value in *T. obturator*. This is consistent with earlier research that did not sample every worker in a colony; isolated individuals are usually less rhythmic than colonies (*3*, *11–13*), and larger groups of workers extracted from colonies tend to be more rhythmic than smaller groups (*13*). For the *L. crassipilis* colony, which had a lower rhythmicity score than the other two colonies, the colony-level rhythmicity fell within the range of the individual-level rhythmicity values. In the case of the ultradian period (the trait which did not show evidence for emergent evolution), we found that the colony-level periods of *T. obturator* and *T. rudis* were close to the mean value of isolated individuals.

### Supplementary Figures

**Figure S1.** The phylogeny of species used in this study, with boxplots showing the walking speed data from isolated individuals. Each data point represents the walking speed of a unique individual during her respective open field assay.

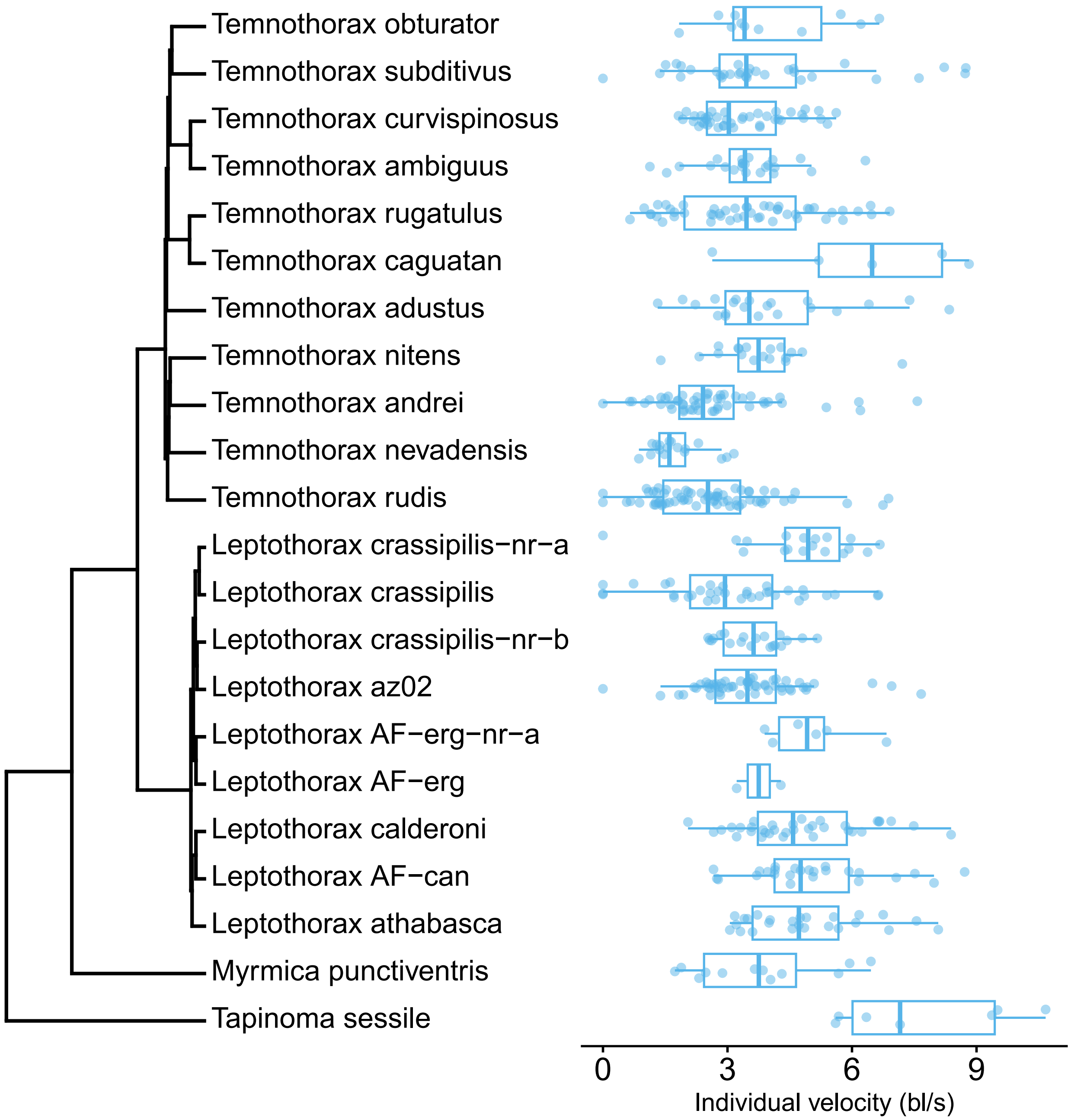

**Figure S2.** (a-b) Scatter plots of the dominant period or rhythmicity of colonies' collective activity vs. the colony size (i.e., approximate number of adult ants inside the nest at the time of recording). Data points are colored according to species. Each data point represents a different time series. Because most colonies were recorded twice, multiple datapoints thus correspond to the same colony measured on different days. The solid lines depict the LME regression for each species. (c-d) Phylomorphospace plots of the dominant period or rhythmicity of species' collective activity vs. the colony size. Each data point represents a different species. The values of these datapoints were the coefficient values obtained from fitting LME models with colony-level period or collective rhythmicity as the response variable and colony size as the fixed effect. Colony ID was used as a random effect in both models.

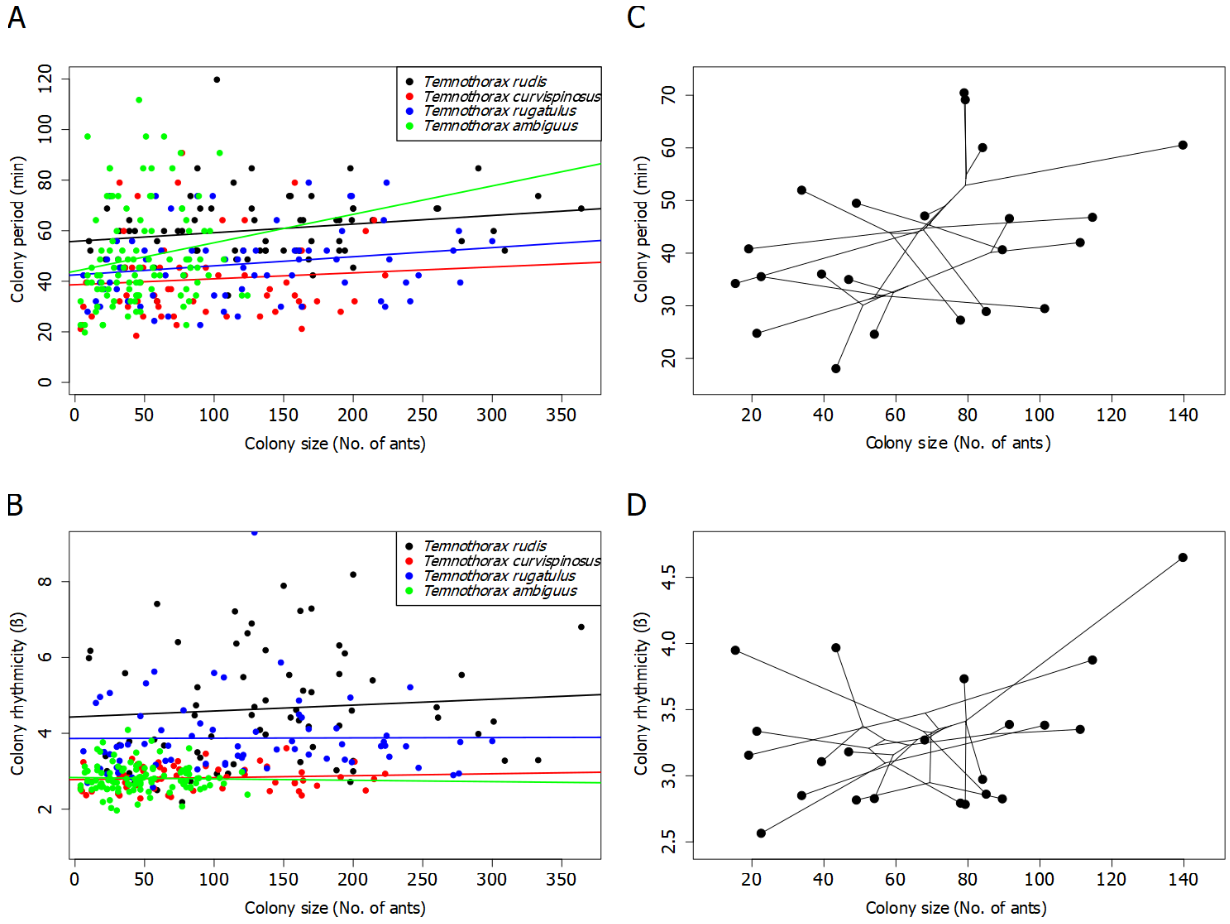

**Figure S3** Histograms of individual-level trait distributions of the constituent ants from the three colonies used in the disassembly experiment. Each column corresponds to one of the three species (*T. rudis*, *T. obturator*, and *L. crassipilis*). The red dotted line in each panel represents the colony-level value for each trait. Specimen images are to scale and were extracted from images from www.antweb.org (casent0005689, casent0104756, and casent0104820).

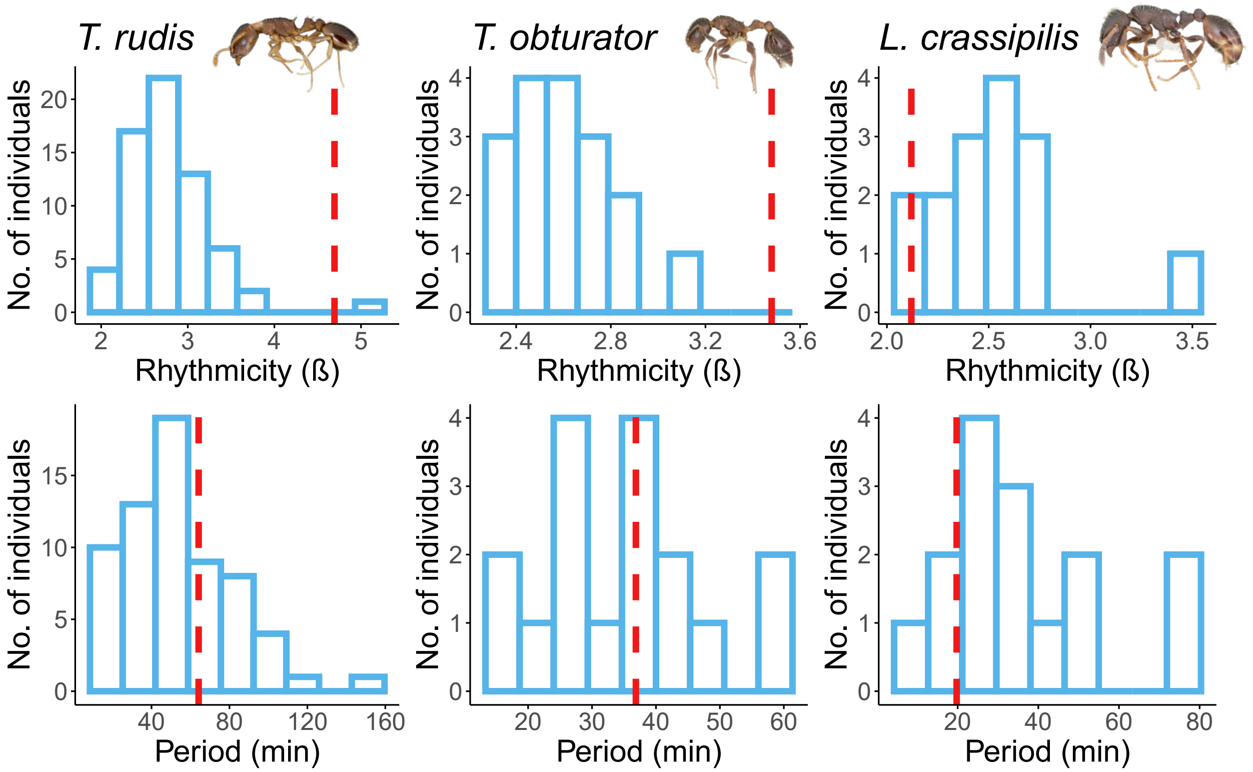
